# Regulation of *CYP94B1* by WRKY33 controls apoplastic barrier formation in the roots leading to salt tolerance

**DOI:** 10.1101/2020.08.10.244608

**Authors:** Pannaga Krishnamurthy, Bhushan Vishal, Wan Jing Ho, Felicia Chien Joo Lok, Felicia Si Min Lee, Prakash P Kumar

## Abstract

Salinity is an environmental stress that causes decline in crop yield. *Avicennia officinalis* and other mangroves have adaptations such as ultrafiltration at the roots aided by apoplastic cell-wall barriers to thrive in saline conditions. We studied a Cytochrome P450 gene, *AoCYP94B1* from *A. officinalis* and its *Arabidopsis* ortholog *AtCYP94B1* that are involved in apoplastic barrier formation, and are induced by 30 minutes of salt treatment in the roots. Heterologous expression of *AoCYP94B1* in *atcyp94b1 Arabidopsis* mutant and wild-type rice conferred increased NaCl tolerance to seedlings by enhancing root suberin deposition. Histochemical staining and GC-MS/MS quantification of suberin precursors confirmed the role of CYP94B1 in suberin biosynthesis. Using chromatin immunoprecipitation, yeast one-hybrid and luciferase assays, we identified AtWRKY33 as the upstream regulator of *AtCYP94B1* in *Arabidopsis*. In addition, *atwrky33* mutants exhibited reduced suberin and salt sensitive phenotypes, which were rescued by expressing *35S::AtCYP94B1* in *atwrky33* mutant. This further confirms that the regulation of *AtCYP94B1* by AtWRKY33 is part of the salt tolerance mechanism, and our findings can help in generating salt tolerant crops.

**One sentence summary:** AtWRKY33 transcription factor regulates *AtCYP94B1* to increase plant salt tolerance by enhanced suberin deposition in the endodermal cells of *Arabidopsis* roots

## Introduction

Salinity is a major environmental stress factor that leads to reduced crop productivity. The progressive increase in soil salinization exacerbates the already damaging effect of steady reduction in the area of arable land worldwide (Parida and Das, 2005; Agarwal et al., 2014). Na^+^ is the major toxic ion found in high saline soils, which imparts osmotic as well as ionic stresses. It is imperative to limit the entry of excess Na^+^ into plant cells in order to maintain proper ion homeostasis, and normal metabolism. Mangroves have evolved various adaptive strategies to flourish under high saline conditions. One of the important adaptations exhibited by most plants, and to a greater extent by mangroves, is ultrafiltration at the roots by the presence of apoplastic barriers in the roots (Scholander, 1968). In an earlier study, we have shown that a salt secretor mangrove, *A. officinalis* restricts 90-95 % salt at the roots due to the presence of enhanced apoplastic barriers (Krishnamurthy et al., 2014).

The main apoplastic diffusion barriers in roots are: epidermis, which is the outermost layer of young roots, endodermis surrounding the vasculature of young roots and peridermis which replaces both epidermis and endodermis in the older roots upon secondary thickening (Nawrath et al., 2013; Wunderling et al., 2018). These apoplastic barriers mainly consisting of Casparian strips (CSs) and suberin lamellae (SL) block the apoplastic and coupled transcellular leakage of ions and water into the xylem, which is the major path of Na^+^ uptake (Yeo et al., 1987; Ma and Peterson, 2003; Krishnamurthy et al., 2011; Kronzucker and Britto, 2011; Schreiber and Franke, 2011; Andersen et al., 2015; Barberon et al., 2016). While CSs are formed as radial wall thickenings, SL are secondary wall thickenings on the inner face of primary cell-walls (Schreiber et al., 1999; Naseer et al., 2012). Chemically, CSs are made up of mainly lignin and SL are made up of suberin and/or lignin depositions (Schreiber et al., 1999; Naseer et al., 2012). Together, these barriers function in biotic and abiotic stress responses (Enstone et al., 2003; Krishnamurthy et al., 2009; Chen et al., 2011; Schreiber and Franke, 2011; Ranathunge et al., 2011a). Suberin is a biopolymer consisting of aliphatic and aromatic domains, with the aliphatic domain contributing mainly to its barrier properties (Kolattukudy, 1984; Schreiber et al., 1999; Ranathunge and Schreiber, 2011b). Suberin biosynthesis is a complex pathway involving elongases, hydroxylases and peroxidases (Bernards et al., 2004; Franke et al., 2005; Hofer et al., 2008; Franke et al., 2009). Cytochrome P450s (CYPs) are one of the largest super-families of peroxidases that are well characterized and known to carry out ω-hydroxylation of the aliphatic constituent of suberin, namely, ω-hydroxy acids (Hofer et al., 2008; Compagnon et al., 2009; Pinot and Beisson, 2011). Most of the CYPs that act on fatty acids belong to *CYP86* and *CYP94* subfamilies (Pinot and Beisson, 2011). Some of the CYPs, such as *CYP86A1*, *CYP94A1*, *CYP94A2* and *CYP94A5* have been identified as ω-hydroxylases (Franke and Schreiber, 2007; Hofer et al., 2008). Although the role of *CYP94B* subfamily in initial ω-oxidation of JA-Ile to 12OH-JA-Ile affecting the JA signaling was known (Koo et al., 2014; Bruckhoff et al., 2016), their role in suberin biosynthesis has not been explored so far and it would be desirable to examine the correlation between *CYP94B* members and root barrier formation.

There is limited information on the molecular mechanisms controlling the genes that regulate suberin biosynthesis. Overexpression of *MYB41* showed increase in suberin biosynthesis as well as expression of some *CYP86* subfamily genes (Kosma et al., 2014). Recently, MYB39 was shown to regulate suberin deposition (Cohen et al., 2020). In addition, the promoter regions of *CYP83* and *CYP71* subfamilies have the W-box, a WRKY transcription factor (TF) binding domain (Xu et al., 2004; Birkenbihl et al., 2017). However, to the best of our knowledge, identities of transcription factor(s) that regulate *CYP94B* subfamily genes is unknown.

In the current study, we have identified and functionally characterized a *CYP94B* subfamily gene, *AoCYP94B1* from *A. officinalis* and its *Arabidopsis* ortholog *AtCYP94B1*. The expression of these genes is induced by salt treatment. We also show that heterologous expression of *AoCYP94B1* increased the salt tolerance and root suberin deposition in *Arabidopsis* and rice seedlings. Histochemical staining was carried out to visualize root suberin, and quantification of suberin precursors in the *atcyp94b1* mutant was done by gas chromatography and mass spectrometry (GC-MS/MS). Using mutant analysis, chromatin immunoprecipitation, yeast one-hybrid, and luciferase assays, we demonstrate that AtWRKY33 regulates *AtCYP94B1*. Additionally, rescue of the reduced suberin and salt sensitive phenotype was demonstrated by expressing *35S::AtCYP94B1* in *atwrky33* mutant. Collectively, the data presented helped to identify the molecular regulatory mechanism involving apoplastic barrier formation through *CYP94B1*, which can be used as an important strategy for generating salt tolerant crops.

## Results

### Identification of *AoCYP94B1*, a *CYP94B* subfamily member from *A. officinalis* as a salt-induced gene

Several cytochrome P450 genes in the *CYP94B* subfamily such as, *AoCYP94B1* and *AoCYP94B3* were identified in our earlier transcriptomic study of *A. officinalis* roots (Krishnamurthy et al., 2017). Since some reports (Benveniste et al., 2006) suggest a role for this subfamily genes in ω-hydroxylation, an important step in suberin biosynthesis, we chose *AoCYP94B1* for further characterization. A phylogenetic tree was constructed based on the derived amino acid sequence of *AoCYP94B1* with other members of this subfamily (Supplemental Figure S1A). Rice OsCYP94B3 and *Arabidopsis* AtCYP94B1 were among the homologs that share high level of sequence similarity with AoCYP94B1. AoCYP94B1 showed 60 % identity and 74 % similarity with AtCYP94B1, and 60 % identity and 71 % similarity with OsCYP94B3. The Cytochrome P450 cysteine heme-iron ligand signature motif was conserved across various plant species (Supplemental Figure S1B).

In *A. officinalis* seedlings without salt treatment, the *AoCYP94B1* transcripts were constitutively expressed in all tissues, but higher level of expression was observed in the leaves and stems compared to roots (Figure 1A). The transcript levels in the roots increased 18-fold with 30 min of NaCl treatment and declined thereafter over 48 h (Figure 1B). In the leaves, 14-fold increase was seen after 4 h of NaCl treatment (Figure 1B). In a parallel exploratory study, the *Arabidopsis* ortholog, *AtCYP94B1* showed somewhat comparable expression pattern in all the tissues tested except for the flowers where it was ~7-fold higher (Figure 1C). The transcript level of *AtCYP94B1* was induced (~4-fold) by 30 min of salt treatment in the roots and remained high up to 6 h. Whereas in leaves, the expression peaked to ~4-fold after 6 h of salt treatment (Figure 1D). Mimicking the qRT-PCR expression profile, the *pAtCYP94B1::GUS* expression was found in all the tissues (Figure 1E, F) and was increased by 30 min upon salt treatment in the roots (Supplemental Figure S2A). While the *pAtCYP94B1::GUS* expression was mainly seen in the stele of the control roots, the expression significantly increased (~3-fold) in salt-treated roots and was mainly found in the endodermis (Figure G and inset to G). Similarly, upon salt treatment, the translational pAtCYP94B1::AtCYP94B1-GFP (Figure 1H, I) fusion localized to endodermal cells where apoplastic barriers are formed.

**Figure 1.**
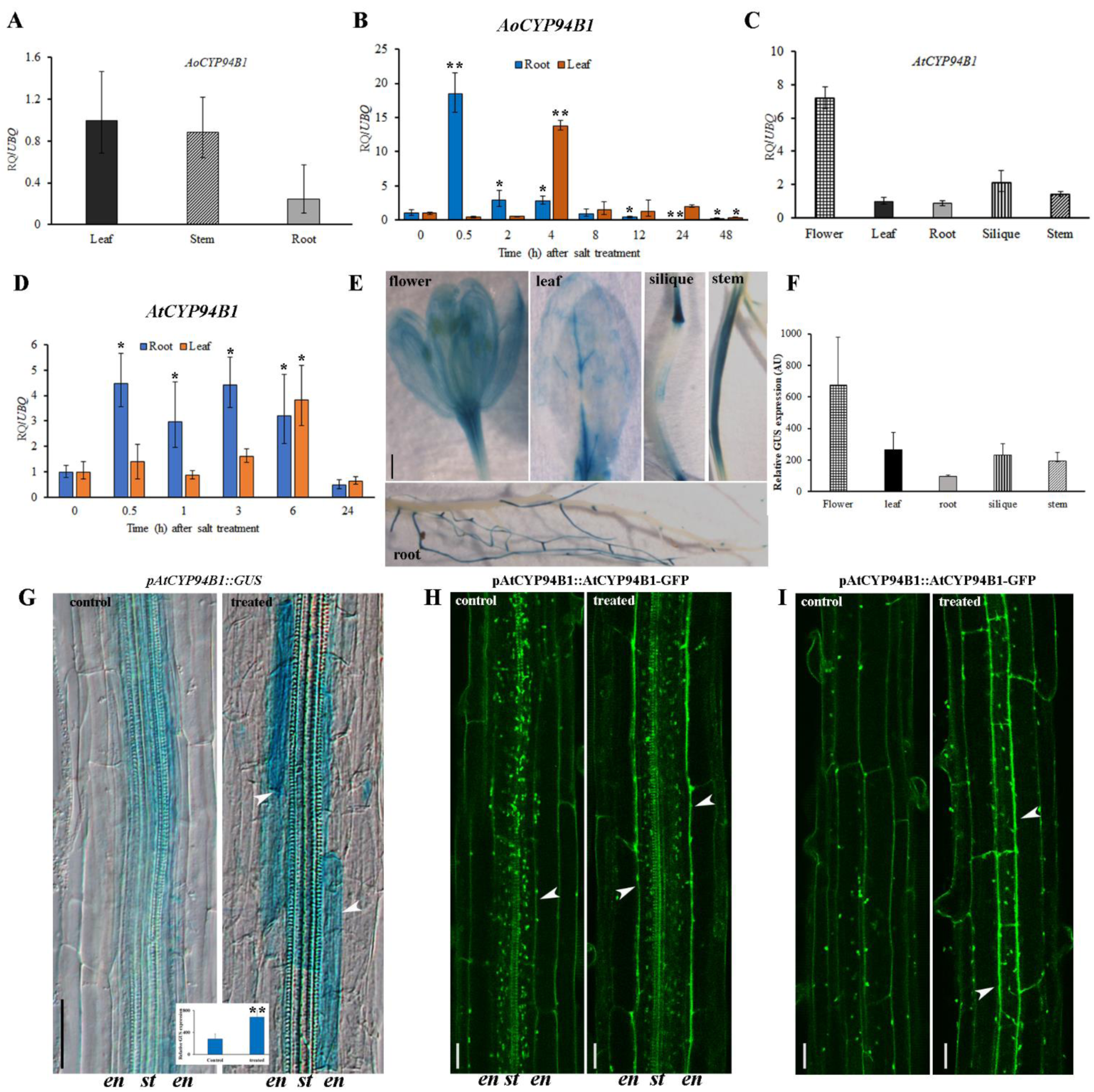
*CYP94B1* is induced by salt stress in both *A. officinalis* and *Arabidopsis*: **(A-B)** Gene expression analyses by qRT-PCR of *AoCYP94B1* in 2-month-old *Avicennia officinalis* plants, (**A**) tissue-specific expression and (**B**) temporal expression in roots and leaves after 500 mM NaCl treatment for varying time periods. (**C**) Tissue-specific expression of *AtCYP94B1* by qRT-PCR in one-week-old *Arabidopsis* seedlings. (**D**) Temporal expression of *AtCYP94B1* in roots and leaves after 50 mM NaCl treatment for varying time periods. Relative expression levels of transcripts with reference to *AtUbiquitin10* and *AoUbiquitin1* transcript levels are plotted in *Arabidopsis* and *A. officinalis*, respectively. The qRT-PCR data represent means ± SD from 3 biological replicates each with 3 technical replicates. (**E**) *pAtCYP94B1::GUS* expression in various tissues of mature plants bearing siliques. Scale bar= 500 µm. (**F**) Relative quantification of *pAtCYP94B1::GUS* expression of E (**G**) Root endodermal cells showing *pAtCYP94B1::GUS* expression in control and salt-treated (50 mM NaCl for 3 h) one-week-old *Arabidopsis* seedlings, scale bar=100 µm. Inset to G; Relative quantification of GUS intensity of G. Data are mean ± SE of three biological replicates, each biological replicate consisting of at least six plants. (**H**) Median and (**I**) surface views of AtCYP94B1-GFP expression in the root endodermal cells of control and salt-treated (50 mM NaCl for 24 h) one-week-old *Arabidopsis* seedlings viewed under confocal microscope, en: endodermis, st: stele. Scale bar=20 µm. Arrowheads in G-I show endodermal cells. Asterisks in all the graphs indicate statistically significant differences (*=*P*< 0.05, **=*P*< 0.01) as measured by Student’s *t*-test between control and the treatments.

### Heterologous expression of *AoCYP94B1* increases salt tolerance in *Arabidopsis* and rice seedlings

In order to functionally characterize the AoCYP94B1, it was heterologously expressed in the *atcyp94b1 Arabidopsis* T-DNA insertional mutant background. There was a reduction in seedling root growth of all the genotypes tested under salt treatment. However, *atcyp94b1* mutants showed about 53% and 76% reduction in root growth upon 50 and 75 mM NaCl treatment, respectively compared to their untreated counterparts (Figure 2A, B). Whereas under similar salt conditions, all the *35S::AoCYP94B1* lines tested grew better than the *atcyp94b1* mutant and WT (34% and 52% growth reduction) with lines 1, 2 and 3 showing 28%, 21%, 17% reduction at 50mM and 43%, 43% and 40% reduction at 75mM, respectively (Figure 2A, B). These data suggest that introduction of *35S::AoCYP94B1* into *atcyp94b1* mutant increased its salt tolerance. In addition, salt sensitivity of 4-week-old plants grown in the soil was checked to see if similar salt response is seen in older plants. Under untreated conditions, there was no difference in the growth of different genotypes. Upon NaCl treatment, *35S::AoCYP94B1* plants displayed better growth compared to the *atcyp94b1* plants (Figure 3A and Supplemental Figure S2B). Yellowing and drying of *atcyp94b1* leaves could be seen and they could not recover to the extent of WT and *35S::AoCYP94B1* lines after the stress was withdrawn. While more than 80% of the WT and *35S::AoCYP94B1* lines showed survival (with more green and healthy leaves) after recovery growth, only 33% of *atcyp94b1* plants could survive the treatment (Figure 3B). There was no significant difference in the leaf area among the genotypes, although a reduction in effective leaf area could be seen due to curling up of leaves upon salt treatment in all the genotypes (Figure 3C). Other growth parameters such as chlorophyll content and FW/DW were measured and found to be generally reduced in all the genotypes upon salt treatment. While *atcyp94b1* mutants showed 3-and 4.5-fold reduction in chlorophyll content and FW/DW ratio respectively, these reductions occurred to a lesser extent (~1.5-fold) in WT and *35S::AoCYP94B1* lines (Figure 3D, E). These observations suggest that introduction of *35S::AoCYP94B1* into *atcyp94b1* mutant rescues its salt sensitive phenotype even in the older plants.

**Figure 2.**
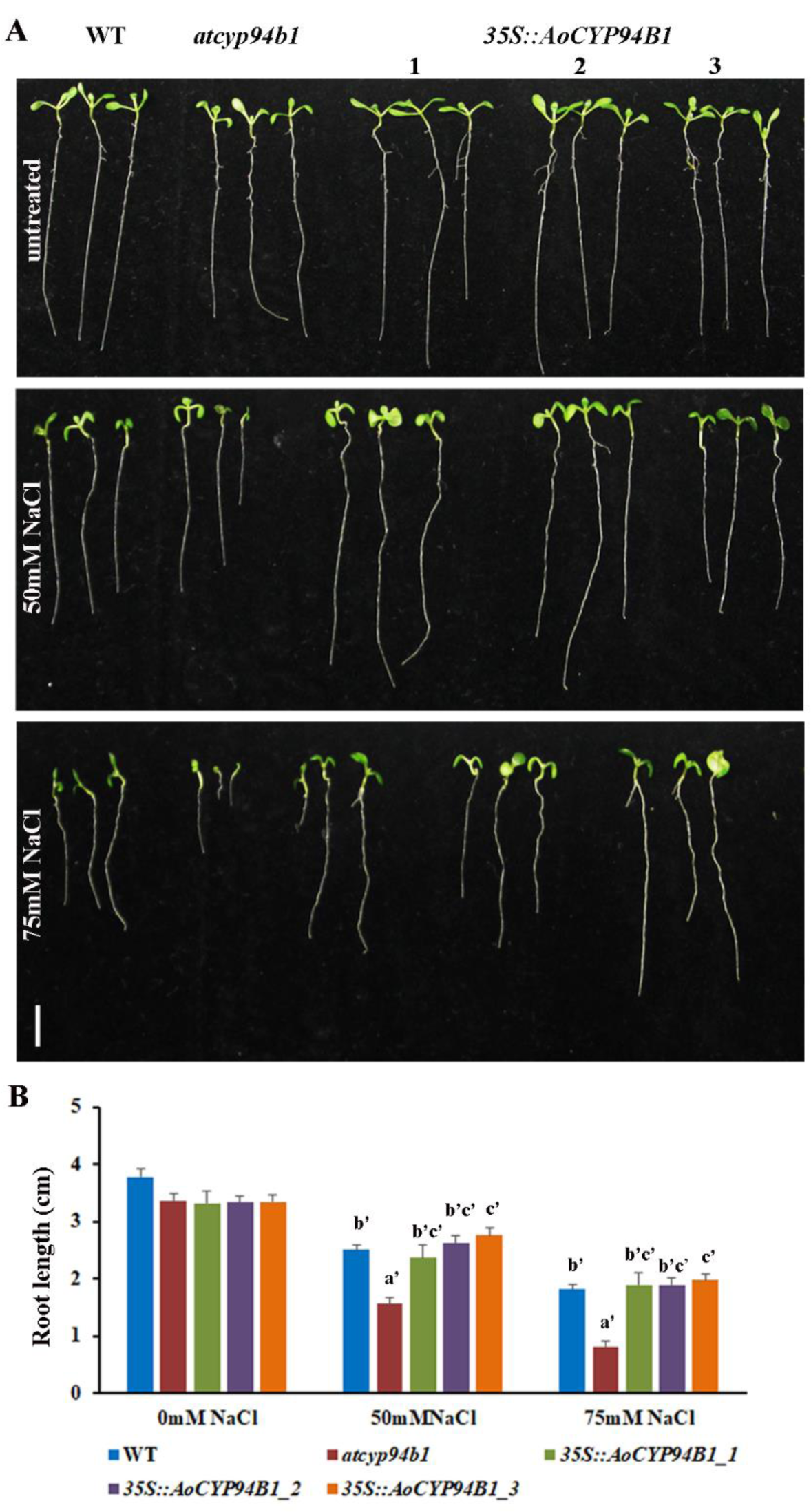
Heterologous expression of *AoCYP94B1* increases salt tolerance in *Arabidopsis* seedlings: (**A**) Comparison of seedling growth among WT, *atcyp94b1* mutant and three independent lines of *35S::AoCYP94B1* heterologously expressed in the mutant background. (**B**) Root growth rates under salt treatment in WT, *atcyp94b1* and *35S::AoCYP94B1* transgenic lines. Surface sterilized and cold stratified seeds were sown on MS agar plates with or without NaCl (50 and 75 mM). Photographs and root length measurements were taken at the end of one week after germination. Data represent mean ± SE of three independent experiments each with at least 15 replicates per experiment. Different letters indicate statistically significant differences between genotypes as determined by the ANOVA employing the Tukey-Kramer posthoc test (*P*<0.01). Same letters indicate no statistical difference between them. Scale bar=10 mm.

**Figure 3.**
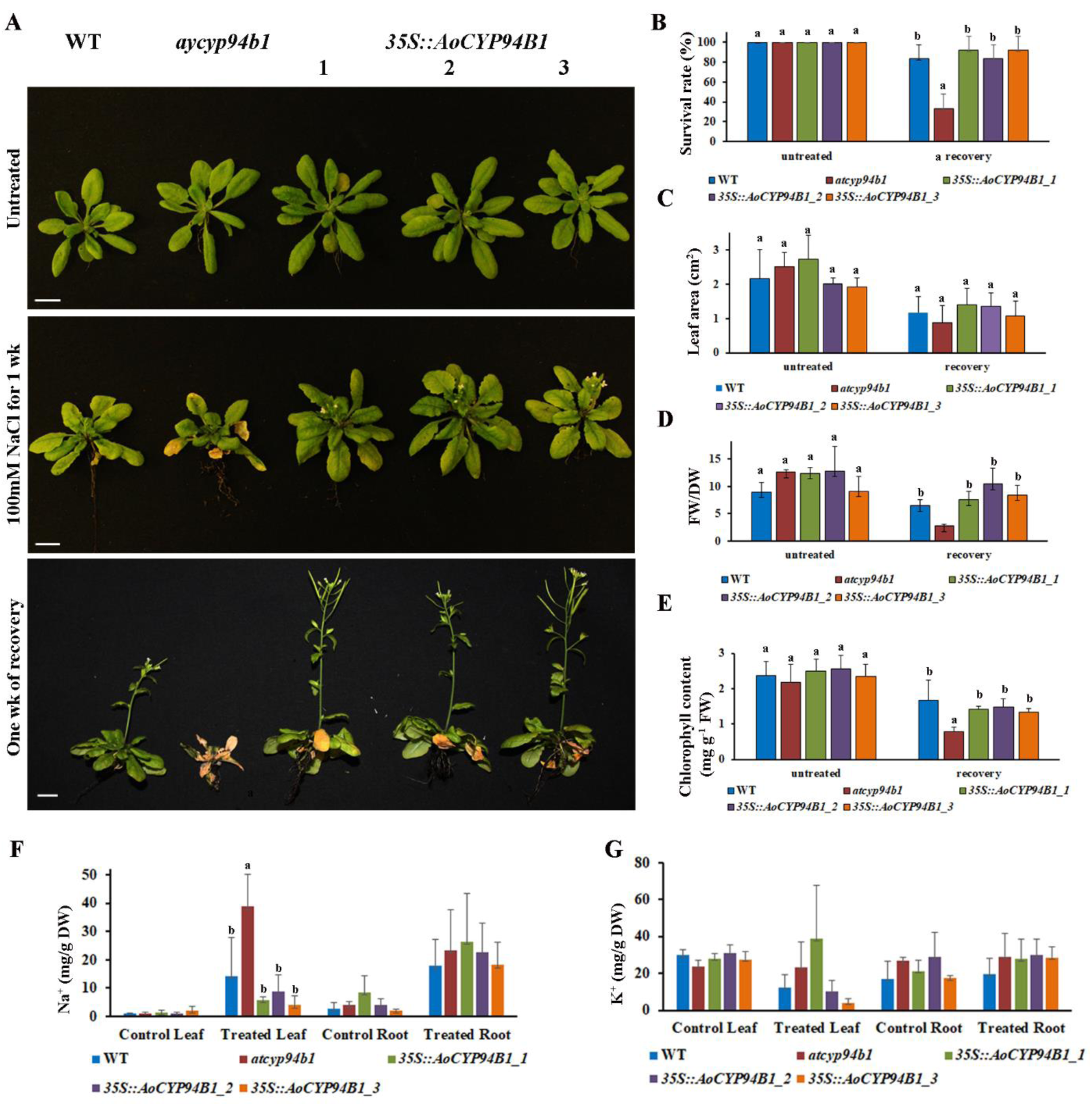
Heterologous expression of *AoCYP94B1* increases salt tolerance and regulates Na^+^ accumulation in *Arabidopsis* plants: (**A-E**) Growth response to salt (100 mM NaCl for 1 week) was monitored in one-month-old, soil grown WT, *atcyp94b1* mutant and three independent lines of *35S::AoCYP94B1* heterologously expressed in the mutant background. (**A**) Growth of the plants shown under untreated condition, under salt-treated condition and after recovery growth in normal water for one week. Scale bar=10 mm. wk; week. Various growth parameters such as (**B**) survival rate (n=12) (**C**) leaf area (n=12) (**D)** FW/DW ratio and (n=12) (**E)** chlorophyll content (n=5) of untreated and recovered *Arabidopsis* plants. (**F**) Total Na^+^ content in the leaves and roots as well as (**G**) total K^+^ content in the leaves and roots of 4-week-old WT, *atcyp94b1* and three *35S::AoCYP94B1* lines. Data are mean ± SE of three biological replicates, each biological replicate consisting of at least 3 plants. Different letters indicate statistically significant differences between genotypes as determined by the ANOVA employing the Tukey-Kramer posthoc test (*P*<0.05). Same letters indicate no statistical difference between them.

We measured the total Na^+^ and K^+^ ion contents in the leaves and roots of WT, mutant and *35S::AoCYP94B1* plants under untreated and salt-treated (100 mM NaCl for 2 days) conditions in order to understand the ion accumulation and distribution. There were no differences in the ion contents among the WT and transgenic lines in the absence of salt treatment. Upon NaCl treatment, the amount of Na^+^ increased from 1 to 38 mg/g DW in the leaves of *atcyp94b1* mutants, while in the *35S::AoCYP94B1* leaves, the amount was significantly lower (6, 8 and 7 mg/g DW in lines 1, 2 and 3, respectively) (Figure 3F). No significant differences were observed in the Na^+^ and K^+^ concentrations within the roots of the different genotypes tested (Figure 3F, G). These data indicate that the *35S::AoCYP94B1* lines efficiently control endogenous Na^+^ accumulation.

Based on the observation that heterologous expression of *35S::AoCYP94B1* in *Arabidopsis* conferred increased salt tolerance, we expressed *pUBI::AoCYP94B1* in rice to examine if a similar increase in salt tolerance could be conferred to the model crop species. There was no difference in the growth of WT and *pUbi::AoCYP94B1* seedlings under untreated conditions (Figure 4A, B). However, the *pUbi::AoCYP94B1* rice seedlings showed significantly higher shoot and root growth than the WT after 3 and 6 days of 100 mM NaCl treatment on MS agar plates (Figure 4C, D). Although growing the young seedlings is convenient on MS medium, we sought to test the rice plants under hydroponic culture conditions normally used to simulate its natural growing environment. On salt treatment for 21 d, the hydroponically-grown one-month-old *pUbi::AoCYP94B1* seedlings showed about 35 % higher survival rate compared to the WT (Figure 4E, F). This further demonstrates that *AoCYP94B1* plays an important role in salt tolerance and could serve as an important candidate for improving salt tolerance of crops.

**Figure 4.**
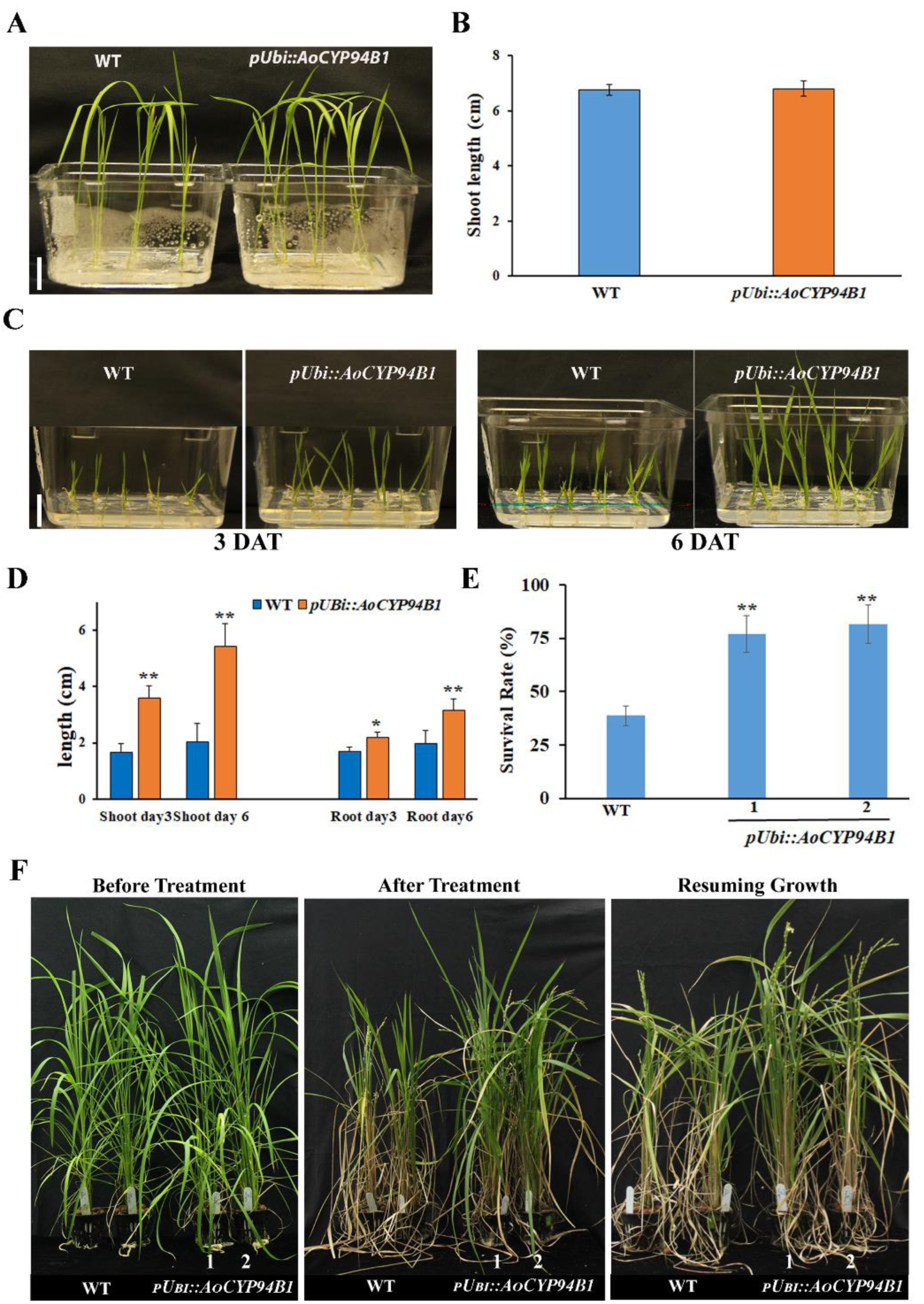
Heterologous expression of *AoCYP94B1* increases salt tolerance in transgenic rice seedlings: **(A)** Phenotype of untreated 2-week-old WT and *pUBI::AoCYP94B1* seedlings **(B)** Shoot length of untreated WT and *pUBI::AoCYP94B1* seedlings **(C)** Phenotype of one-week-old WT and *pUBI::AoCYP94B1* seedlings after 100 mM NaCl treatment. **(D)** Shoot and root lengths of WT and *pUBI::AoCYP94B1* seedlings after three and 6 days of salt treatment. **(E)** 4-week-old WT and *pUBI::AoCYP94B1* plants grown in hydroponics, before salt treatment, after 21 days of 100 mM NaCl treatment and an additional 10 days of recovery growth **(F)** Survival rates of WT and *pUBI::AoCYP94B1* plants after salt treatment and recovery growth. Data in (B, D and F) are mean ± SD of three independent experiments each with at least 15 seedlings per experiment. Asterisks indicate statistically significant differences (\*\**P*< 0.01) between *pUBI::AoCYP94B1* line and WT as measured by Student’s *t*-test. Scale bar=1 cm, DAT; days after treatment.

In order to gain further insights into the underlying molecular mode of action, we chose to work with the *Arabidopsis* ortholog. This would permit more detailed biochemical and molecular genetic analyses to be performed, which would not be feasible with *Avicennia*, a perennial tree species that is not amenable to genetic transformation.

### *AtCYP94B1* increases salt tolerance as well as suberin lamellae (SL) formation in *Arabidopsis* roots

Similar to the *AoCYP94B1* heterologous expression lines, *pAtCYP94B1::AtCYP94B1* complementation lines showed better seedling root growth compared to the *atcyp94b1* mutants when subjected to 50 and 75 mM NaCl treatment (Figure 5A-D). The salt sensitivity phenotype of *atcyp94b1* mutants was rescued in the *pAtCYP94B1::AtCYP94B1* complementation lines. Analysis was carried out in three *pAtCYP94B1::AtCYP94B1* complementation lines and two representative lines are shown in Figure 5A-C. However, mannitol treatment, as an alternate abiotic stress, did not cause significant changes to the seedling root growth in any of these genotypes (Supplemental Figure S3).

**Figure 5.**
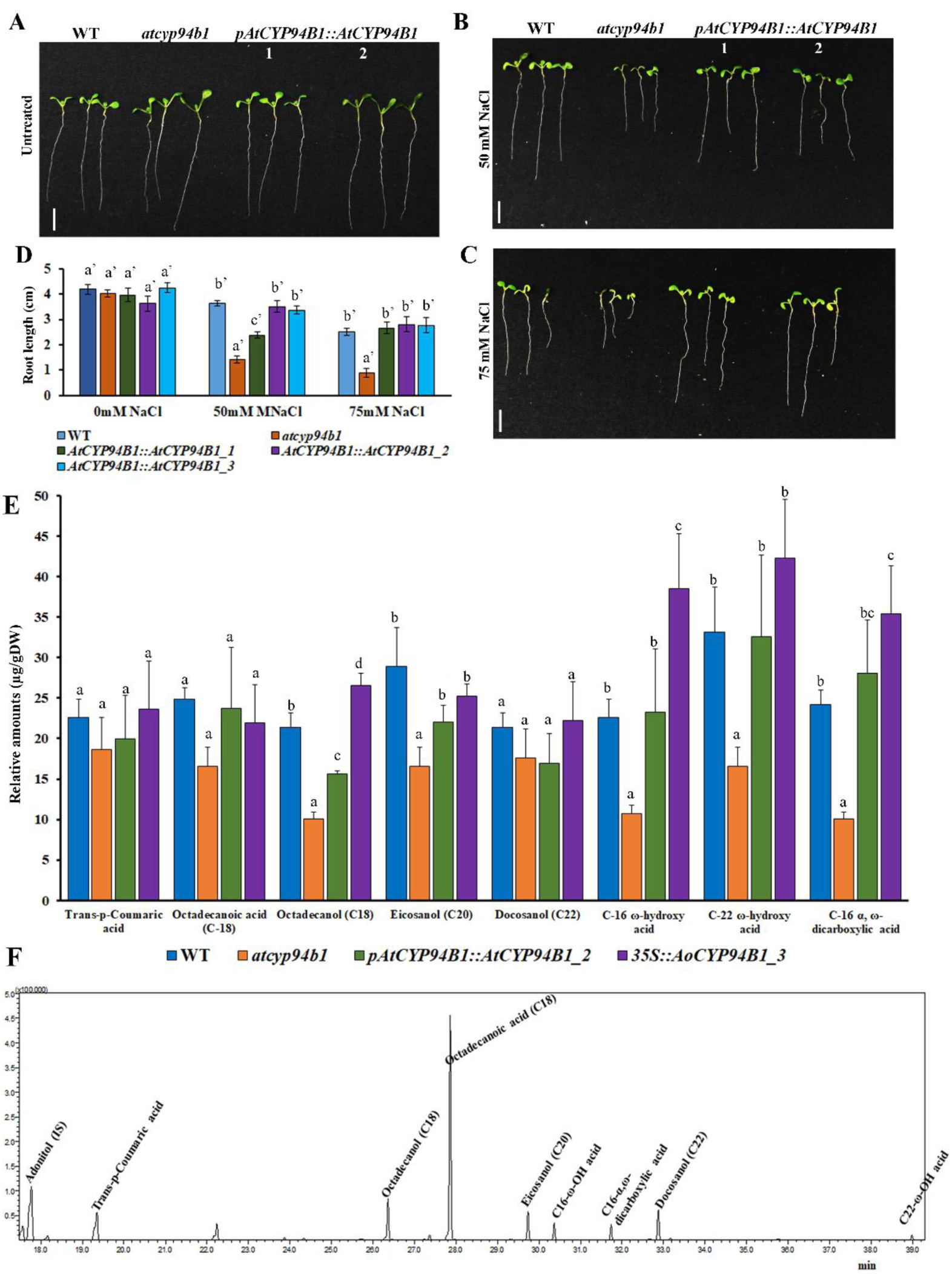
Complementation of *atcyp94b1* with *Arabidopsis AtCYP94B1* increases salt tolerance and suberin levels in *Arabidopsis* roots: (**A-C**) Comparison of seedling growth among WT, *atcyp94b1* mutant and two independent complementation lines of *AtCYP94B1::AtCYP94B1* in mutant background. (**D**) Root growth rates under salt treatment in WT, *atcyp94b1* and *AtCYP94B1::AtCYP94B1* complementation lines. Surface sterilized and cold stratified seeds were sown on MS agar plates with or without NaCl (50 and 75 mM). Photographs and root length measurements were taken at the end of one week after germination. Data are mean ± SE of at least 15 biological replicates. Different letters indicate statistically significant differences between genotypes as determined by the ANOVA employing the Tukey-Kramer posthoc test (*P*<0.01). Same letters indicate no statistical difference between them. Scale bar=10 mm. **(E)** Suberin monomer composition in the seedling roots of 4-week-old WT, *atcyp94b1*, *pAtCYP94B1::AtCYP94B1* and *35S::AoCYP94B1* were quantified using GC-MS/MS analysis. Data are mean ± SD of three independent biological replicates each with 4-5 plants. Different letters indicate statistically significant differences between genotypes as determined by the ANOVA employing the Tukey-Kramer posthoc test (*P*<0.05). Same letters indicate no statistical difference between them. **(F)** Chromatogram (multiple reaction monitoring) for the standard suberin monomers and internal standard (adonitol).

In view of the suggested role for *CYP94B* family genes in suberin biosynthesis, combined with our observation of reduced Na^+^ accumulation in the shoots of *35S::AoCYP94B1* lines, we carried out GC-MS/MS quantification of several aliphatic components of the root suberin monomers in WT, *atcyp94b1* mutant, *pAtCYP94B1::AtCYP94B1* complementation lines and *35S::AoCYP94B1* heterologous expression lines. The *atcyp94b1* mutant showed a significant reduction in the amount of ω-hydroxy acids and α, ω-dicarboxylic acid compared to the other genotypes tested (Figure 5E). No significant differences in the amounts of p-coumaric acid, C-18 octadecanoic acid and C-22 docosanol was found between the genotypes tested. While *atcyp94b1* mutant showed ~50 % reduction in the amounts of alcohols (C-18 octadecanol and C-20 eicosanol), ω-hydroxy acids (C-16 and C-22) and C-16 α, ω-dicarboxylic acid (Figure 5E), the amounts were restored to the WT levels in *pAtCYP94B1::AtCYP94B1* complementation lines. The increase seen in *35S::AoCYP94B1* lines was higher compared to that of the WT, which could be either due to the strength of 35S promoter or because *AoCYP94B1* functions much more efficiently. Similarly, significantly higher amounts of C-16 ω-hydroxy acid and C-16 α, ω-dicarboxylic acid were present in *35S::AoCYP94B1* line compared to WT indicating a role for CYP94B1 in their biosynthesis. All the standards quantified for this study are shown in the chromatogram in Figure 5F.

Further, to visualize the altered suberin deposition in the roots, we carried out root histochemical studies in WT, *atcyp94b1* mutant, *pAtCYP94B1::AtCYP94B1* complementation lines and *35S::AoCYP94B1* heterologous expression lines. There was no difference in the deposition of root CSs among the genotypes (Figure 6B). However, there was a significant reduction in the deposition of suberin in the endodermal cell walls of *atcyp94b1* compared to WT, *pAtCYP94B1::AtCYP94B1* and *35S::AoCYP94B1* roots. While SL was found in ~70 % of the endodermal cells in WT, *pAtCYP94B1::AtCYP94B1* and *35S::AoCYP94B1*, only 20 % of the cells exhibited SL in *atcyp94b1* mutant (Figure 6C-E). In the patchy zones of *35S::AoCYP94B1_1* roots, a significantly higher number of endodermal cells with suberin were seen (Figure 6C, E). Furthermore, we examined the uptake of fluorescein diacetate (FDA), which was used as a tracer to check the barrier properties of SL previously (Barberon et al., 2016). In the undifferentiated (CSs not formed) and non-suberized (well-formed CSs) zones of the roots of all the genotypes checked, after 1 min of incubation, FDA could enter all the endodermis (100 %) as well as the pericycle cells (Supplemental Figure S4A-D). However, in the suberized zones of the roots, FDA could penetrate only ~10 % of endodermal cells in WT, *pAtCYP94B1::AtCYP94B1* and *35S::AoCYP94B1*, while it entered ~80 % of *atcyp94b1* endodermal cells (Figure 5F, G). In addition, to check if the enhanced salt tolerance in rice is brought about by a similar mechanism of action, namely, increased SL deposition as seen in *Arabidopsis*, we examined the salt-treated roots of WT and *pUbi::AoCYP94B1* rice lines. In the apical regions, SL were visible in a higher number of endodermal cells of *pUbi::AoCYP94B1* than in the WT. In the mid regions, SL were clearly visible in *pUbi::AoCYP94B1* while several endodermal cells lacked SL in the WT. In the basal regions, prominent SL were present in all the endodermal cells of *pUbi::AoCYP94B1*, while many passage cells without SL were evident in the WT (Supplemental Figure S5A vs. B). The SL deposition showed a similar trend in the exodermal layer as in the endodermis (Supplemental Figure S5C vs. D). These results not only suggest that *AtCYP94B1* has a critical role in the formation of SL as the apoplastic barrier leading to salt tolerance, but also that *AoCYP94B1* could function similar to its *Arabidopsis* ortholog. Therefore, the use of *AtCYP94B1* for further understanding of its molecular regulatory mechanism can be justified.

**Figure 6.**
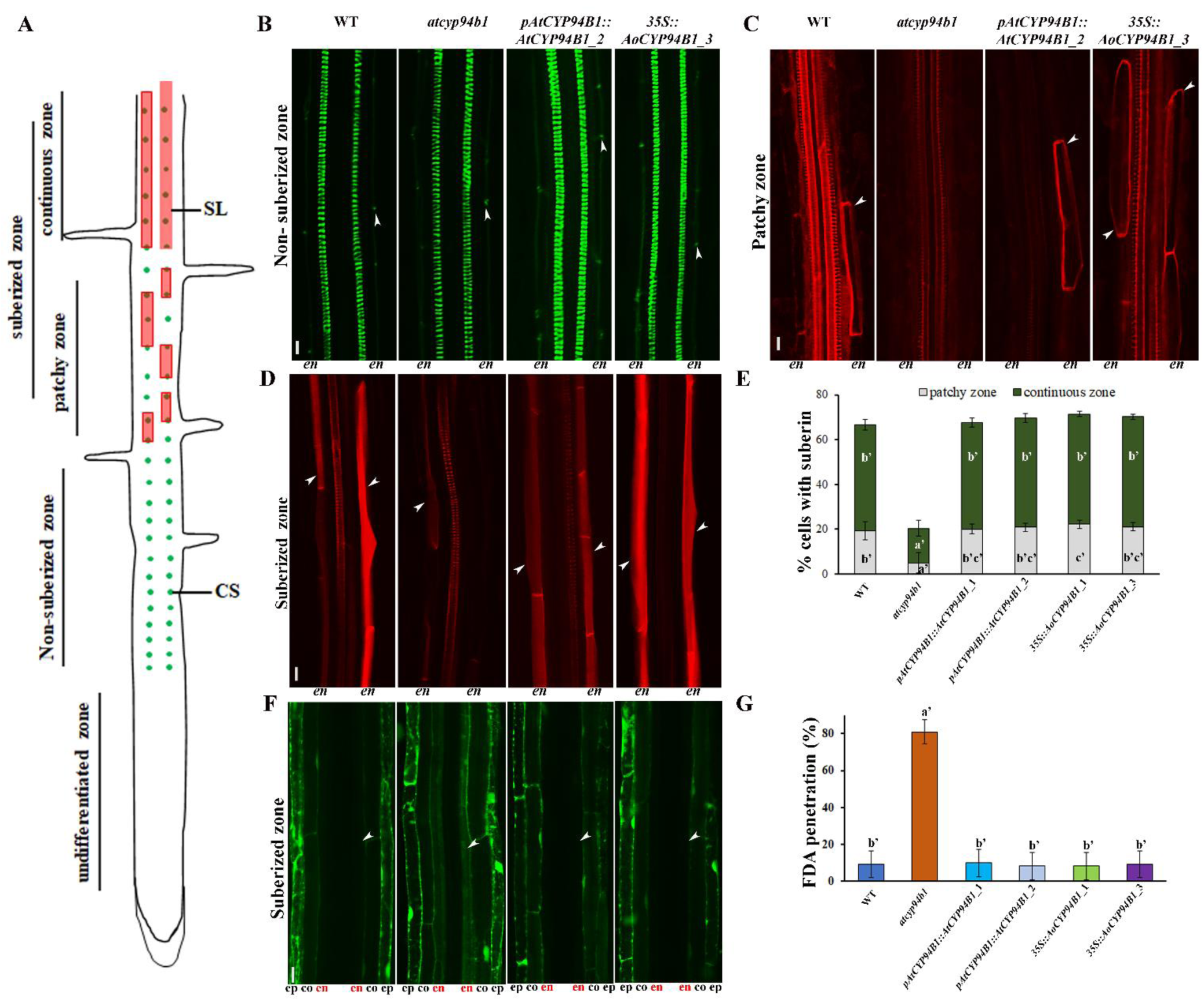
CYP94B1 is involved in apoplastic barrier (SL) formation in *Arabidopsis* roots: For root anatomical studies, one-week-old *Arabidopsis* seedlings grown on MS agar plates were used. Images were taken from similar parts of the WT, *atcyp94b1*, *pAtCYP94B1::AtCYP94B1* and *35S::AoCYP94B1* stained roots. Seedlings were stained with Auramin O to visualize CSs, and with Nile Red to view SL. Suberin patterns were counted as described in materials and methods. **(A)** Schematic of endodermal differentiation (adapted from (Barberon et al., 2016)). Three different zones are shown: undifferentiated, non-suberized, and suberized zone (patchy and continuous zones are distinguished). (**B**) Representative images showing CSs in the endodermis of non-suberized zones of roots. (**C, D**) Images showing SL deposition in the endodermal cells of patchy and continuous suberized zones of roots. (**E**) Percentage of endodermal cells with SL in the suberized zones. n=10 seedlings. (**F**) FDA penetration after 1 min in the suberized root zones of WT, *atcyp94b1*, *pAtCYP94B1::AtCYP94B1* and *35S::AoCYP94B1*. **(G)** Percentage of endodermal cells with FDA penetration in the suberized zone. ep: epidermis, co: cortex, en: endodermis (in red). Arrowheads indicate the location of CS and SL, except in F where arrowheads show the endodermis, Scale bar=10 µm. Different letters indicate statistically significant differences between genotypes as determined by the ANOVA employing the Tukey-Kramer posthoc test (*P*<0.01). Same letters indicate no statistical difference between them.

### Identification of AtWRKY33 transcription factor as the upstream regulator of AtCYP94B1

We sought to identify the upstream regulator of *AtCYP94B1* in *Arabidopsis* after establishing the fact that *AoCYP94B1* and *AtCYP94B1* function in a similar manner to regulate root apoplastic barrier formation leading to salt tolerance. Analysis of the 5’-upstream region of *AtCYP94B1* showed various abiotic stress-related *cis*-elements such as WRKY, MYB and MYC transcription factor binding domains, and especially, an enrichment of WRKY binding domains (Supplemental Figure S6A). Coincidentally, in our earlier transcriptomic study, WRKYs (e.g., AoWRKY6, AoWRKY9, AoWRKY33) were one of the major groups of TFs upregulated upon salt treatment in the roots of *A. officinalis* (Krishnamurthy et al., 2017). Because the role of WRKY33 in salt tolerance of plants has emerged in several studies, we selected AtWRKY33 which shares high sequence similarity to AoWRKY33, (Supplemental Figures S7 A, B) for our further study. Also, we used WRKY6 and WRKY9 that belong to Group I in ChIP and Y1H assays to ensure that the interaction and regulation is specific to WRKY33 (Group I WRKY). We carried out qRT-PCR analysis to check if *AtWRKY33* was induced by salt treatment in a similar manner as *AtCYP94B1*. Under untreated control conditions, we observed that *AtWRKY33* expression was comparable across tissue samples (Figure 7A). In contrast, as was seen for *AtCYP94B1* earlier, expression levels of *AtWRKY33* increased by 30 min of salt treatment in the roots and remained high (4-fold after 3 h) up to 6 h (Figure 7B). Similarly, in leaves, *AtWRKY33* expression increased 5-fold with 30 min of salt treatment (Figure 7B). Furthermore, similar to the qRT-PCR expression profile, the *pAtWRKY33::GUS* expression was seen in all the tissues in untreated seedlings (Supplemental Figure S8) and increased upon 50 mM NaCl treatment in the roots (Figure 7C, D). Surprisingly, the *pAtWRKY33::GUS* was mainly expressed in salt-treated *Arabidopsis* root endodermal cells (Figure 7E) similar to *AtCYP94B1* expression.

**Figure 7.**
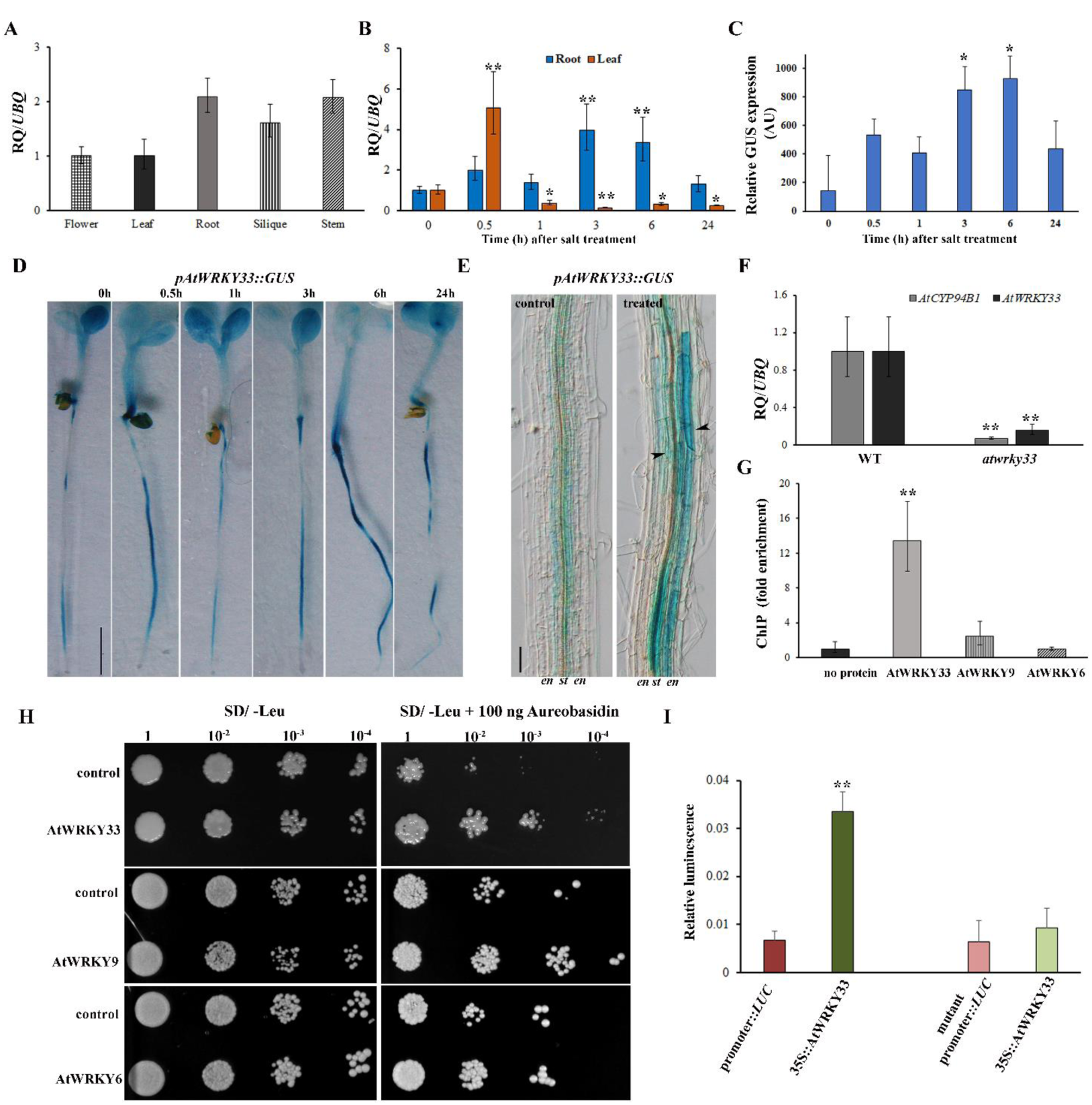
AtWRKY33 transcription factor acts as an upstream regulator of *AtCYP94B1*: (**A-B**) Gene expression analyses by qRT-PCR of *AtWRKY33* in one-week-old *Arabidopsis* seedlings. (**A**) Tissue-specific expression, (**B**) temporal expression in roots and leaves after 50 mM NaCl treatment for varying time periods. (**C**) Relative quantification and (**D**) *pAtWRKY33::GUS* expression analysis in one-week-old seedling roots upon 50 mM NaCl treatment. Scale bar=500 µm. Asterisks indicate statistically significant differences (*=*P*< 0.05, **=*P*< 0.01) between 0 h and other time points as measured by Student’s *t*-test. (**E**) Root endodermal cells showing *pAtWRKY33::GUS* expression in control and salt-treated (50 mM NaCl for 6 h) one-week-old *Arabidopsis* seedlings, scale bar=100 µm. Arrowheads show endodermal cells. (**F**) Suppression in the transcript levels of *AtCYP94B1* in *atwrky33* T-DNA insertional mutant roots compared to WT. Asterisks indicate statistically significant differences (**=*P*< 0.01) between WT and *atwrky33* mutant as measured by Student’s *t*-test. (**G**) Chromatin immunoprecipitation (ChIP)-qPCR of HA-tagged AtWRKY33 in *Arabidopsis* protoplasts, AtWRKY6 and AtWRKY9 were used as negative controls. Fold change in the enrichment of promoter fragments compared to no protein control are plotted. qRT-PCR data represent means ± SD from 3 biological replicates each with 3 technical replicates. Asterisks indicate statistically significant differences (**=*P*< 0.01) between no protein control and AtWRKY33 as measured by Student’s *t*-test. (**H**) Yeast one-hybrid assay showing regulation of *AtCYP94B1* by AtWRKY33. AtWRKY6 and AtWRKY9 were used as additional controls. The representative growth status of yeast cells is shown on SD/-Leu agar medium with or without 100 ng of aureobasidin A. Numbers on the top of each photograph indicate relative densities of the cells 4 days post-inoculation. (**I**) Luciferase assay was carried out using the mesophyll protoplasts obtained from the leaves of 4-week-old *atwrky33* mutants. The *pAtCYP94B1::LUC* was used as the control and *35S::AtWRKY33* was used as the test. *AtCYP94B1* promoter fragment with mutated WRKY binding sites was used as additional control. Firefly luciferase activity was normalized to Renilla luciferase activity and plotted. Data represent mean ± SD of four independent biological replicates each with three technical replicates. Asterisks indicate statistically significant differences (**=*P*< 0.01) as measured by Student’s *t*-test between the control and the test.

To experimentally validate whether AtWRKY33 regulates *AtCYP94B1*, the expression level of *AtCYP94B1* was quantified in *atwrky33* mutants. *AtCYP94B1* transcript levels decreased by 14-fold in *atwrky33* mutants (Figure 7F). In addition, ChIP-qPCR analysis was performed to check for WRKY interaction with *AtCYP94B1* promoter fragment. Consistent with the presence of putative WRKY-binding *cis*-elements, over tenfold enrichment of *AtCYP94B1* promoter fragment was observed in AtWRKY33-HA pulldown samples (Figure 7G). AtWRKY6-HA and AtWRKY9-HA pulldown was also carried out to check if the interaction was specific to AtWRKY33, and we found that there was no significant enrichment of AtCYP94B1 promoter fragments in these pulldown samples. We independently verified the interaction of AtWRKY33, AtWRKY6 and AtWRKY9 with the promoter fragment of *AtCYP94B1* using the Y1HGold system (Clontech USA). After introduction of *pGADT7 AtWRKY33, AtWRKY6* and *AtWRKY9* plasmids into the Y1HGold cells harboring *AtCYP94B1* promoter fragment, AtWRKY33 grew better than its control in the presence of Aureobasidin A (100 ng ml^-1^), indicating an interaction between AtWRKY33 and the promoter of *AtCYP94B1* (Figure 7H). While AtWRKY6 did not show any better growth compared to the control, there was very weak interaction with AtWRKY9 (Figure 7H). Additionally, luciferase assay using *atwrky33 Arabidopsis* mutant protoplasts was carried out to check the *in vivo* transcriptional activation of *AtCYP94B1* promoter by AtWRKY33. Protoplasts transfected with *pAtCYP94B1::LUC* along with *35S::AtWRKY33* showed ~3-fold higher luminescence compared to the ones transfected with the control, *pAtCYP94B1::LUC* (Figure 7I). The mutant *pAtCYP94B1::LUC* (with two WRKY binding sites mutated) showed only ~1.5-fold higher luminescence compared to the control indicating that the mutation in the TF binding sites indeed affects the promoter activity. Collectively, these results show that AtWRKY33 TF acts as the upstream regulator of *AtCYP94B1* gene.

If the identified WRKY33 is indeed the upstream regulator of *AtCYP94B1 in vivo*, root apoplastic barrier deposition in the *atwrky33* mutants should be impaired and expression of *35S::AtCYP94B1* in *atwrky33* mutant should rescue this phenotype. There were no visible differences in the formation of CSs in the *atwrky33* mutants compared to WT and *35S::AtCYP94B1 atwrky33* roots (Figure 8A). However, suberin deposition was reduced in the roots of *atwrky33* compared to the WT and *35S::AtCYP94B1 atwrky33* (Figure 8B). While ~70 % of WT and 60 % of *35S::AtCYP94B1 atwrky33* endodermal cells showed SL deposition, only 25 % of the corresponding *atwrky33* cells exhibited SL deposition (Figure 8C). Further, the *atwrky33* mutant seedlings showed salt sensitivity similar to that shown by *atcyp94b1* mutants. However, this sensitivity was rescued when *35S::AtCYP94B1* was expressed in *atwrky33* mutant background (Figure 8D). These results strongly support our proposed working model where AtWRKY33 regulates root apoplastic barrier formation via *AtCYP94B1* to confer enhanced salt tolerance in plants (Figure 8E).

**Figure 8.**
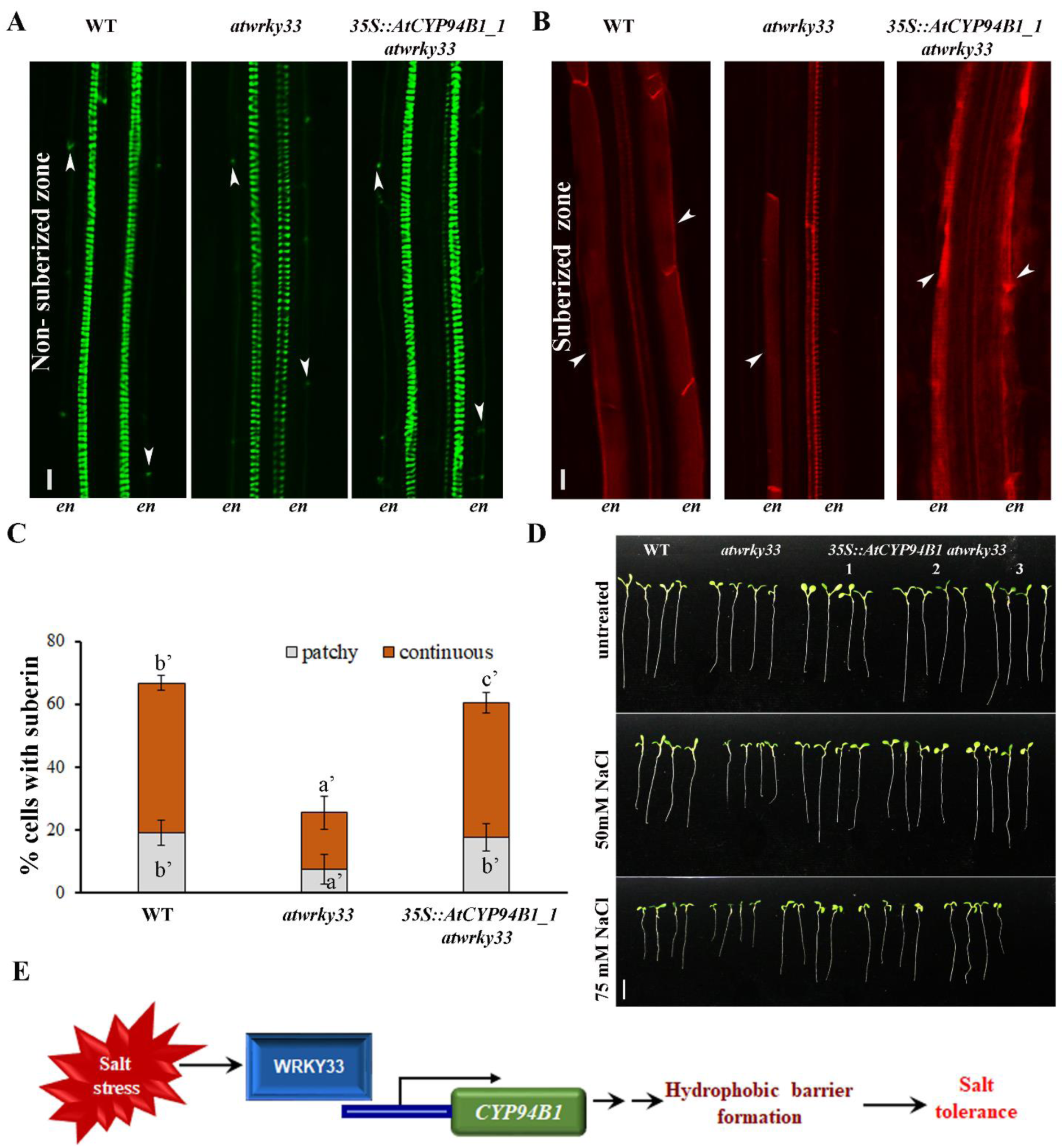
AtWRKY33 regulates apoplastic barrier formation via *AtCYP94B1*: For root anatomical studies, one-week-old WT, *atwrky33* mutants and *35S::AtCYP94B1 atwrky33* (in the *atwrky33* mutant background) *Arabidopsis* seedlings grown on MS agar plates were used. Images of stained roots were taken from the same regions for all the genotypes. For visualizing CSs, seedlings were stained with Auramin O, while they were stained with Nile Red to view SL. Suberin patterns were counted as described in materials and methods. (**A**) Representative images showing Casparian strip development in the non-suberized endodermal cells of all three genotypes. (**B**) SL deposition in the suberized endodermal cells of all three genotypes. (**C**) Percentage of endodermal cells with SL in the suberized zones of the roots. Arrowheads indicate the location of CS and SL, n=10 seedlings, Scale bar=10 µm. Different letters indicate statistically significant differences between genotypes as determined by the ANOVA employing the Tukey-Kramer posthoc test (*P*<0.01). Same letters indicate no statistical difference between them. (**D**) Comparison of seedling growth among WT, *atwrky33* mutant and three independent lines of *35S::AtCYP94B1 atwrky33* ectopic expression lines. Surface sterilized and cold stratified seeds were sown on MS agar plates with or without NaCl (50 and 75 mM). Photographs were taken at the end of one week after germination. Scale bar=10 mm. en: endodermis. (**E**) Proposed model based on our data showing regulation of root apoplastic barrier formation by WRKY33 through controlling *CYP94B1* leading to salt tolerance.

## Discussion

It is imperative for researchers to understand the mechanisms underlying salt tolerance and generate salt tolerant crops in order to meet the increasing demand for food to support the predicted population growth. Various studies have shown that apoplastic barrier (mainly CSs and SL) deposition in the root endodermis and exodermis is critical in order to prevent unwanted loading of ions into the xylem (Krishnamurthy et al., 2011; Schreiber and Franke, 2011; Nawrath et al., 2013; Graca, 2015; Barberon, 2017; Kreszies et al., 2018). Although it is known that mangroves possess highly efficient apoplastic barrier deposition (Krishnamurthy et al., 2014), the underlying molecular mechanism was not understood previously. The potential to learn from such adaptive mechanisms to devise strategies for crop improvement has been highlighted, but that is yet to be accomplished. The present study represents a successful example of discovering and applying such mechanistic knowledge.

To understand the role of CYP94B1 in salt stress response, we have used three plant species (*A. officinalis*, *Arabidopsis* and rice) of varying ages. Our earlier transcriptomic study involving *A. officinalis* which led to the identification of *AoCYP94B1*, was carried out using 2-month-old seedlings treated with 500 mM NaCl. Therefore, similar conditions were used for *A. officinalis* in the current study. Experiments in *Arabidopsis* were carried out using the young seedlings (one-week-old) and older (4-week-old) plants in order to understand their response to salt in two developmental stages. While 50 mM NaCl was used for most of the studies with younger seedlings as this did not damage the roots, 100 mM NaCl was used to challenge the older plants. Similarly, two developmental stages (one-week-old and 4-week-old) of rice plants were used for our studies.

Our findings have highlighted the role of *AtCYP94B1* from *Arabidopsis* in salinity tolerance response. This gene was identified as the ortholog of *AoCYP94B1* based on our studies with the mangrove tree, *A. officinalis*, which suggests that they may play similar roles in the two species. Under untreated conditions, the expression of *AtCYP94B1* was the lowest in roots while it was predominant in flowers (Figure 1D). Similar expression profile of *AtCYP94B1* was reported in earlier studies (Bruckhoff et al., 2016; Widemann et al., 2016). However, according to BAR eFP Browser, the highest expression is found in petioles of mature leaves. Also, *AtCYP94B1* gene family was shown to regulate flowering time but not the fertility (Bruckhoff et al., 2016), while overexpression of *AtCYP94B3* led to partial loss of male sterility in *Arabidopsis* (Koo et al., 2011). However, in our study, ~4-fold upregulation of *AtCYP94B1* seen in the roots upon salt treatment (Figure 1D) along with its expression and localization to the endodermis (Figure 1G-I) clearly indicates its key function in root endodermis under salt stress. While *AoCYP94B1* showed highest expression at 0.5 h after salt treatment, *AtCYP94B1* expression remained high from 0.5 h to 6 h. At this point, we are not sure if this difference is a reflection of inherent differences between two species or due to other reasons.

Earlier studies have suggested that CYP94B family genes, including AtCYP94B1, play a role in sequential ω-oxidation of JA-Ile to 12OH-JA-Ile (Koo et al., 2014; Aubert et al., 2015; Lunde et al., 2019) particularly after flower opening (Bruckhoff et al., 2016; Widemann et al., 2016). The authors also had highlighted that a role for this CYP94 subfamily members in metabolism of other substrates cannot be dismissed. Furthermore, in an earlier study, *Arabidopsis* CYP94B1, expressed in yeast was shown to carry out ω-hydroxylation of fatty acids with chain lengths of C12-C18 (Benveniste et al., 2006). Accordingly, our observation that the reduction in the concentration of aliphatic suberin monomers (C-16 and C-22 ω-hydroxyacids) in the roots of *atcyp94b1* mutants compared to the WT, complementation and heterologous expression lines (Figure 5E) confirms the role of CYP94B1 in biosynthesis of aliphatic suberin monomers, leading to the formation of apoplastic barriers in the roots. The induction of *CYP94B1* gene expression by salt treatment in *Arabidopsis* and *A. offiicnalis* (Figure1), coupled with the observation of reduced apoplastic barrier formation in the root endodermis (Figure 6) as well as salt sensitivity in the *atcyp94b1* mutant (Figures 2–4) collectively show that *AtCYP94B1* plays a role in salinity response. The fact that the salt sensitive phenotype along with reduced amount of suberin in the endodermis of *atcyp94b1* mutant are rescued in the transgenic *Arabidopsis* lines (*35S::AoCYP94B1* and *pAtCYP94B1::AtCYP94B1*) further confirm the role of CYP94B1 in salt tolerance via suberin deposition. It is also clear from our data that the suberin lamellae in the endodermal cell walls are functional in blocking the apoplastic flow (Figure 6F) and thereby limiting the uptake of excess Na^+^ into the shoots (Figure 3F). Several prior studies have shown similar increases in the concentration of aliphatic suberin monomers in response to salt stress which limits the root apoplastic bypass flow (Ranathunge et al., 2005; Ranathunge et al., 2008; Krishnamurthy et al., 2009; Krishnamurthy et al., 2011; Ranathunge et al., 2011a; Krishnamurthy et al., 2014). The increased salt tolerance exhibited by the heterologous expression lines (*35S::AoCYP94B1*) of rice (Figure 4) and *Arabidopsis* (Figure 2, 3) was directly correlated with increased apoplastic barrier formation in the roots (Figures 5, 6, Supplemental Figure S5), suggesting that the mangrove *AoCYP94B1* could be used to confer enhanced salt tolerance to crop plants. It has also been shown that JA-Ile degradation / oxidation occurs under salt stress (Hazman et al., 2019) and this leads to increased salinity tolerance in rice (Kurotani et al., 2015). Therefore, the role of this pathway in contributing to salt tolerance in our overexpression lines cannot be dismissed and demands further studies to understand the relationship between JA catabolism and suberin biosynthesis under salt stress.

Suberin deposition not only occurs in response to abiotic stresses such as salinity, drought (Ranathunge et al., 2011c; Franke et al., 2012), but also to biotic stresses where it serves to block pathogen entry through the cell walls (Ranathunge et al., 2008). Despite their pivotal role in conferring tolerance to multiple stresses, information on molecular regulation of apoplastic barrier formation is still scarce. So far, some of the MYB TFs have been shown to regulate CS development (Kosma et al., 2014; Kamiya et al., 2015; Li et al., 2018) and suberin biosynthesis (Gou et al., 2017) by regulating *CYP86* subfamily genes. To the best of our knowledge, no prior report on regulation of *CYP94B* subfamily genes exists. Salt-mediated co-induction of *AtWRKY33* (Figure. 7B-D) along with *AtCYP94B1* (Figure 1D) and the endodermal expression of both *pAtCYP94B1::GUS* and *pATWRKY33::GUS* in the salt-treated roots (Figures 1, 7) suggests that they both play a related role under salt treatment. This, combined with enrichment of *AtCYP94B1* promoter fragments in our ChIP-qPCR, yeast one-hybrid analysis and luciferase assay (Figure 7) clearly show that AtWRKY33 is the upstream regulator of *AtCYP94B1*. WRKY TFs are known to regulate biotic (Bakshi and Oelmuller, 2014; Sarris et al., 2015) and abiotic stress responses. (He et al., 2016; Liang et al., 2017; Bai et al., 2018). Several transcriptomic and microarray studies have shown their response to salt and drought stresses (Mahalingam et al., 2003; Narusaka et al., 2004; Krishnamurthy et al., 2011; Okay et al., 2014; Song et al., 2016). Additionally, overexpression of WRKY25, 33, 41 and 83 caused enhanced salt tolerance (Jiang and Deyholos, 2009; Chu et al., 2015; Wu et al., 2017), while many others (WRKY46, 54, 68, 70) play a role in drought tolerance (Chen et al., 2017). However, the mechanism by which these WRKY TFs confer stress tolerance is not known. Our observations of reduced deposition of SL in the endodermal cells of *atwrky33* mutants mimicking *atcyp94b1* mutant phenotype along with the increase in SL deposition in the *35S::AtCYP94B1 atwrky33* lines (Figure 8) further confirm the regulatory role of AtWRKY33 transcription factor on *AtCYP94B1*. Hence, we propose a working model where WRKY33 regulates root apoplastic barrier formation by controlling *CYP94B1* leading to increased salt tolerance (Figure 8E).

In conclusion, our study reveals a part of an important molecular regulatory mechanism of suberin deposition that involves the control of *AtCYP94B1* by AtWRKY33 transcription factor, leading to increased salt tolerance of *Arabidopsis* seedlings. We further showed that heterologous expression of *AoCYP94B1* in both *Arabidopsis* and rice seedlings confers salt tolerance by the same mode of action. Therefore, our study opens new avenues for engineering salt tolerant crop plants.

## Materials and Methods

### Cloning and generation of transgenic *Arabidopsis* and rice lines

Full-length coding sequence of *AoCYP94B1* (Unigene99608_All) was obtained from our earlier transcriptomic study of *A. officinalis* roots (Krishnamurthy et al., 2017). Wild-type (WT) *Arabidopsis thaliana*, ecotype Columbia-0 along with the T-DNA insertional mutants, *atcyp94b1* (SALK_129672) and *atwrky33* (SALK_064436) were purchased from the Arabidopsis Biological Resource Center (ABRC) (http://www.abrc.osu.edu) (Alonso et al., 2003). Position of T-DNA insertion sites for *atcyp94b1* and at*wrky33* mutants are shown in Supplemental Figures S9 and S10, respectively. Genomic DNA from the mutants was extracted as described previously (Dellaporta et al., 1983). Plants homozygous for the T-DNA insertion were selected by genotyping with primers designed using the T-DNA primer design tool (http://signal.salk.edu/tdnaprimers.2.html). Also, qRT-PCR was carried out to check the suppression of the tagged gene expression in mutants (Supplemental Figures S9, S10). Seeds were collected from only those mutants that showed more than 70 % reduction in *AtCYP94B1* as well as *AtWRKY33* expression (Supplemental Figures S9C and S10C). For heterologous expression of *AoCYP94B1* in *Arabidopsis*, coding sequence (CDS) of *AoCYP94B1* gene was amplified and cloned into pGreen binary vector under 35S promoter. For construction of translational fusion with GFP and complementation lines of *AtCYP94B1*, the full-length cassette including promoter, introns and exons of *AtCYP94B1* amplified from the genomic DNA of *Arabidopsis* was cloned into pGreen-GFP and pGreen binary vectors, respectively. For promoter GUS expression analysis, 1kb upstream sequences of *AtCYP94B1* and *AtWRKY33* were amplified from genomic DNA and cloned into pGreen-GUS. All the constructs generated were individually electroporated into *Agrobacterium tumefaciens* strain GV3101:pMP90 and introduced into *Arabidopsis* by the floral dip method (Clough and Bent, 1998). Basta-resistant T1 transgenic plants were selected and introduced gene expression was confirmed by genotyping PCR and qRT-PCR analysis (Supplemental Figure S9D). T3 generation plants were used for all the experiments. For chromatin immunoprecipitation (ChIP) assay, coding sequences of *AtWRKY33* was cloned into pGreen binary vector with hemagglutinin (HA) fusion tag. For heterologous expression in rice, coding sequence of *AoCYP94B1* gene was amplified and cloned with compatible restriction into binary vector pCAMBIA-1300 under corn *UBIQUITIN* promoter. The constructs were introduced into rice by *Agrobacterium*-mediated transformation (Toki et al., 2006). Homozygous transgenic lines (3:1 segregation ratio on hygromycin selection) at the T2 generation were selected for further analysis. All the primers used in the study are listed in Supplemental Table 1.

### Plant materials and growth conditions

The propagules of *Avicennia officinalis* L. (*A. officinalis*) were collected during fruiting seasons from the mangrove swamps in Singapore (Berlayer Creek and Sungei Buloh Wetland Reserve). The seedlings were maintained in NaCl-free conditions by growing in potting mixture (Far East Flora, Singapore), until they reached the four-node stage (~2 months) in a greenhouse (25–35o C, 60–90 % relative humidity; 12 h photoperiod), after which they were carefully transferred to pots containing sand and were allowed to adapt for two days by watering with half-strength Hoagland’s solution. The plants were then treated with half-strength Hoagland’s solution containing 500 mM NaCl for varying time periods (0 h, 0.5 h, 1 h, 2 h, 4 h, 8 h, 12 h, 24 h and 48 h).

For growth on Murashige-Skoog (MS) agar plates, *Arabidopsis* seeds of required genotypes were surface sterilized and cold stratified at 4 °C for 3 days, then the seeds were sown on MS agar plate and germinated at 22 °C under continuous light. One-week-old seedlings were carefully removed from the plate and subjected to salt (50 mM NaCl) treatment. The plant tissues were collected at various time periods (0 h, 0.5 h, 1 h, 3 h, 6 h and 24 h) and frozen in liquid nitrogen for total RNA isolation. For histochemical GUS expression analysis, One-week-old seedlings were treated with 50 mM NaCl for various time periods (0 h, 1 h, 3 h, 6 h and 24 h). For root length studies, the sterilized and cold stratified seeds were sown on MS agar plate with and without NaCl and the root lengths were measured and photographed One week after germination. For salt treatment in older plants, 4-week-old plants were treated with 100 mM NaCl for one week. The pots were flushed with water twice to remove the soil-bound NaCl followed by a recovery growth in NaCl-free water for one week. Survival rate, chlorophyll contents, FW/DW ratio and leaf area were measured from untreated and recovered plants. In addition, untreated plants bearing mature siliques were collected for qRT-PCR analysis and GUS staining.

Rice seeds of WT (*Oryza sativa* subsp. *japonica* cv. Nipponbare) and heterologous expression lines (*pUbi::AoCYP94B1*) were first germinated on plain MS or selection medium (MS+hygromycin) and then transferred to MS agar plates with and without NaCl (100 mM NaCl). After one week of growth in this medium, seedlings were photographed and the shoot length and root lengths were measured. WT and the transgenic lines generated were grown hydroponically (Yoshida et al., 1971) for salt treatment of older seedlings. Four-week-old seedlings were subjected to salt treatment (100 mM NaCl) for 21 days. After treatment, the rice seedlings were transferred back to NaCl-free hydroponic solution for recovery. Survival rates were calculated after 10 days of recovery and are shown as the percentage of seedlings that were alive. Plants that did not show any indication of recovery (no green shoots) were counted as dead.

### *In silico* analysis

The NCBI database was used as a search engine for nucleotide and protein sequences. Expasy tool (https://web.expasy.org/translate/) was used to translate the CDS sequences to amino acid sequences and multAlin (http://multalin.toulouse.inra.fr/multalin/) was used to align the amino acid sequences. Phylogenetic analysis was carried out using http://www.phylogeny.fr/. Primers for qRT-PCR were designed using NCBI (https://www.ncbi.nlm.nih.gov/tools/primer-blast/).

### RNA isolation and Quantitative real-time PCR (qRT-PCR) analysis

RNA was isolated from various plant samples (*A. officinalis*, *Arabidopsis* and rice) using TRIzol reagent (Thermo Fisher) following the manufacturer’s instructions. An aliquot of this RNA (1 µg) was used to synthesize cDNA using Maxima first strand cDNA synthesis kit for qRT-PCR (Thermo Fisher) following the manufacturer’s instructions. For genotyping of mutants and the heterologous expression lines, RNA was extracted from leaves of 4-week-old seedlings. The qRT-PCR to check expression of transcript levels was performed using StepOne*^TM^* Real-Time PCR machine (Applied Biosystems, Foster City, CA, USA) with the following programme: 20 s at 95 °C followed by 40 cycles of 03 s at 95 °C and 30 s at 60 °C. The SYBR Fast ABI Prism PCR kit from KAPA (Biosystems, Wilmington, MA, USA) was used for qRT-PCR analysis. The reaction mixture consisted of 5.2 μl master mix (provided in the kit), 0.2 μM each of forward (FW) and reverse (RV) primers, 3.4 μl nuclease-free water, and 1 μl sample cDNA template for a final volume of 10 μl. All of the data were analyzed using the StepOne*^TM^* Software (v2.1, ABI). The primers used for the qRT-PCR analysis are listed in Supplemental Table 1. Gene expression levels were calculated based on ∆∆CT values and represented as relative expression levels (fold change) to constitutively expressed internal controls, *AtUbiquitin 10* and *AoUbiquitin 1* for *Arabidopsis* and *A. officinalis*, respectively.

### Chlorophyll estimation

Chlorophyll concentrations were determined spectrophotometrically using 100 mg FW of untreated and recovered (from NaCl treatment) leaf material ground in 2 ml of acetone 80% (v/v). After complete extraction, the mixture was filtered and the volume adjusted to 5 ml with cold acetone. The absorbance of the extract was read at 663 and 645 nm and pigment concentrations were calculated as described previously (Arnon, 1949). The data represented are mean ± SD of 4 biological replicates each with single plants.

### Estimation of total ion concentration (Na^+^ and K^+^) from plants

Control and salt-treated 4-week-old *Arabidopsis* seedlings were harvested and rinsed briefly with distilled water to remove surface contaminating Na^+^. Pool of three to four plants was taken as one replicate, and three to four independent replicates were used to generate the mean values reported. Leaves and roots from plants were separated and left to dry at 50 °C for 2 days. The dried tissue was ground into a powder in liquid nitrogen, and acid digestion and ion estimation were carried out as described earlier (Krishnamurthy et al., 2014). The amounts of ions estimated are presented as mg / gDW of plant sample.

### Histochemical GUS staining

Transgenic *Arabidopsis* seedlings containing *pAtCYP94B1::GUS* and *pAtWRKY33::GUS* fusion constructs were treated as described above. GUS histochemical staining was performed by vacuum infiltrating the seedlings immersed in GUS staining solution [0.1 M sodium phosphate buffer pH 7.0, 10 M EDTA, 0.1% Triton X-100, 2 M 5-bromo-4-chloro-3-indolyl glucuronide (X-Gluc)] for 5 min followed by overnight incubation in the dark at 37 °C without shaking. Staining solution was removed and several washes with 50 % ethanol were performed until the chlorophyll was bleached and tissues cleared. The images of stained whole seedlings with various salt treatments were recorded using a stereo microscope (NIKON SMZ1500) and other *pAtCYP94B1::GUS* images were taken using LEICA CTR5000 DIC microscope with a Nikon DS-Ril camera. GUS-stained images presented here represent the typical results of at least six independent plants for each treatment. GUS expression was quantified based on the relative intensities of blue coloration using ImageJ software. Data presented are mean ± SD of three biological replicates, each biological replicate consisting of at least six plants.

### Chemical analysis of suberin in the root

Isolation and depolymerization of suberized root cell walls from 4-week-old *Arabidopsis* seedlings were carried out as described previously (Franke et al., 2005; Hofer et al., 2008). Briefly, the samples were depolymerized by transesterification with 2 ml 1M MeOH/HCl for 2 h at 80 °C followed by addition of NaCl/H2O. 10 μg of adonitol was added as internal standard and aliphatic monomers were extracted (three times in 0.5 ml) in hexane. The combined organic phase was dried using CentriVap Cold Traps (Labconco) and derivatized using bis-(N,N,-trimethylsilyl)-tri-fluoroacetamide (BSTFA; Sigma) as described previously (Franke et al., 2005; Hofer et al., 2008). Monomers were identified from their EI-MS spectra (75 eV, m/z 50-700) after capillary GC (DB-5MS, 30 m x 0.32 mm, 0.1 μm, on column injection at 50 °C, oven 2 min at 50 °C, 10 °C min^-1^ to 150 °C, 1 min at 150 °C, 5 °C min^-1^ to 310 °C, 30 min at 310 °C and He carrier gas with 2 ml min^-1^) on Shimadzu gas chromatograph combined with a quadrupole mass selective detector (GCMS-TQ80). Quantitative analysis of suberin monomers was carried out based on normalization to the internal standard. Three biological replicates, each consisting of 4-5 plants from all the genotypes tested, were used for the analysis. Suberin amounts are presented as µg/g DW of the sample.

### Histochemical staining and microscopy

For root histochemical studies, one-week-old *Arabidopsis* seedlings grown on MS agar were used. Seedlings were stained with Auramin O and Nile Red to visualize CS and SL, respectively following the methods described earlier (Ursache et al., 2018). For CS development, cells from the first CS appearance to the first elongated cell were checked. As described previously, (Barberon et al., 2016) suberin patterns were counted from the hypocotyl junction to the onset of endodermal cell elongation (Onset of elongation is the zone where an endodermal cell length is clearly more than twice its width). The roots were divided into various zones (Figure 6A) such as undifferentiated zone (young part of the root with no CS and SL), non-suberized zone (only CS, no SL) and suberized zone (patchy and continuous SL). For FDA transport assay, seedlings were incubated for 1 min in 0.5 x MS FDA (5 µg ml-1), rinsed, and observed using a confocal laser scanning microscope (FV3000, Olympus). Excitation and detection parameters were set as follows: Auramin O 488 nm, 505-530 nm; Nile Red 561nm, 600-650 nm; FDA 488 nm, 620-640 nm. Images were taken from at least 10 *Arabidopsis* seedlings of each genotype tested for all the analyses.

For microscopy of rice roots, freehand cross-sections were prepared from the adventitious roots of salt-treated rice plants grown in hydroponics. Roots of ~100 mm length were taken from the hydroponically grown, salt-treated plants and were sectioned at varying lengths from the root tip: apical (0–20 mm), mid-(20–50 mm) and basal (50–80 mm). To check for SL deposition, sections were stained for 1 h with Fluorol Yellow 088 (Brundrett et al., 1991). Stained sections were viewed under a confocal laser scanning microscope (FV3000, Olympus) with excitation at 514 nm and detection at 520–550 nm and DAPI filter (excitation at 405 nm, detection at 420–460 nm). Root images shown represent the typical results of at least six independent rice plants.

### Chromatin Immunoprecipitation (ChIP) using *Arabidopsis* protoplasts

Mesophyll protoplasts were isolated from leaves of 4-week-old WT *Arabidopsis* (Col-0) plants and transfected according to the protocol described earlier (Yoo et al., 2007) with minor modifications. For each transfection, 8–15 μg of purified plasmid DNA (*35S::AtWRKY33*, *35S::AtWRKY6* and *35S::AtWRKY9*) was used. The three WRKYs were chosen to ensure that the interaction is specific to WRKY33 (Group I WRKY) and distinct from WRKY6 and WRKY9 that belong to Group I. Polyethylene glycol (PEG)–calcium chloride transfection solution used was as follows: 25 % PEG, 0.4 M mannitol, and 150 mM CaCl_2_. The transfected protoplasts were incubated for 20 h at room temperature and fixed with formaldehyde. Protoplasts transfected with empty vectors were treated as the negative control. Anti-HA monoclonal antibody (Santa Cruz Biotechnology) bound to Protein-A agarose beads (Sigma) were used to immunoprecipitate the genomic DNA fragments. ChIP-qPCR analysis was carried out to check for promoter fragment enrichment in the final eluted chromatin from the ChIP experiment. Fold change in the enrichment of promoter fragments compared to the control were plotted. Results are based on data from three independent biological replicates each with at least three technical replicates.

### Luciferase assay using *Arabidopsis* protoplasts

Mesophyll protoplasts were isolated from leaves of 4-week-old *atwrky33* mutant seedlings and luciferase assay was carried out as described earlier (Iwata et al., 2011). 1kb upstream sequence of *AtCYP94B1* was cloned into pGreen II-0800-LUC vector to generate the reporter. The vector with the *pAtCYP94B1* promoter (*pATCYP94B1::LUC*) was used as reference control (reporter) while *35S::AtWRKY33* was used as effector. Two WRKY binding sites in the *AtCYP94B1* promoter fragment were mutated (TTGAC to TTacC) (Supplemental Figure S6B) by site directed mutagenesis and the mutant promoter was cloned into pGreen II-0800-LUC vector and used as an additional control. The luciferase assay was carried out using the Dual-Luciferase^®^ Reporter Assay System (Promega) following the manufacturer’s instructions. The luminescence was measured using the GloMax discover (Promega). Firefly luciferase activity was normalized to Renilla luciferase activity. Data shown were taken from four independent biological replicates each with three technical replicates.

### Yeast One-hybrid assays (Y1H)

Y1H assays were performed using a Matchmaker™ Gold Y1H System (Clontech, USA) according to the manufacturer’s instructions. The promoter fragment, 2kb upstream of *AtCYP94B1* was cloned into the pAbAi vector upstream of *AUR1-C* reporter gene. Coding sequence of *AtWRKY33*, *AtWRKY6* and *AtWRKY9* were cloned into the pGADT7-AD vector. The primers used for cloning are listed in Supplemental Table 1. The strains were then allowed to grow for 2–3 days at 30 °C to assess DNA–protein interactions.

### Statistical analysis

Data presented are the mean values ± SE / SD. Binary comparisons of data were statistically analyzed by Student’s *t*-test (*P* < 0.05 and *P* < 0.01). For multiple comparisons between wild type, mutant and transgenic lines, one-way analysis of variance (ANOVA) was performed and Tukey-Kramer posthoc test was subsequently used as a multiple comparison procedure (*P* < 0.05 and *P* < 0.01).

## Acknowledgements

This research grant was supported by the Singapore National Research Foundation under its Environment and Water Research Programme and administered by PUB, Singapore's National Water Agency, Singapore, NRF-EWI-IRIS (R-706-000-040-279). We thank the NParks Singapore for granting us permission to collect the mangrove samples from Berlayer Creek and Sungei Buloh Wetland Reserves (NP/RP 12-002-1 & NP/RP 12-002-2).

## Supplemental information

**Supplemental Figure 1:** AoCYP94B1 is highly similar to other plant CYP94B subfamily

**Supplemental Figure 2:** *pAtCYP94B1::GUS* expression and growth of 4-week-old WT, *atcyp94b1* and *35S::AoCYP94B1 Arabidopsis* plants

**Supplemental Figure 3:** WT, *atcyp94b1*, *35S::AoCYP94B1* and *AtCYP94B1::AtCYP94B1* complementation lines of *Arabidopsis* responded similarly to mannitol treatment

**Supplemental Figure 4:** FDA penetration into endodermal cells show functionality of suberin

**Supplemental Figure 5:** Heterologous expression of *AoCYP94B1* increases deposition of apoplastic barrier (SL) in the roots of transgenic rice seedlings

**Supplemental Figure 6:** Promoter analysis of *AtCYP94B1*

**Supplemental Figure 7:** *A. officinalis* WRKY33 shares high sequence similarity with *Arabidopsis* WRKY33

**Supplemental Figure 8:** Tissue specific expression of *pAtWRKY33::GUS*

**Supplemental Figure 9:** Details of *atcyp94b1* mutant and heterologous expression lines **Supplemental Figure 10:** Details of *Arabidopsis wrky33* mutant

**Supplemental Table 1:** Details of primer sequences used in the study

## Supplemental Figures

**Supplemental Fig. 1: AoCYP94B1 is highly similar to other plant CYP94B subfamily:** (**A**) A phylogenetic tree was constructed using the deduced amino acid sequences of *AoCYP94B1* along with *Manihot esculenta* (XP_021615193.1), *Theobroma cacao* (EOY22465.1), *Gossypium hirsutum* (XP_016708735.1), *Vitis vinifera* (XP_002279981.1), *Citrus clementina* (XP_006440103.1), *Arabidopsis thaliana* (NP_201150.1), *Capsicum chinensis* (PHU16576.1), *Capsicum baccatum* (PHT53723.1), *Solanum lycopersicum* (XP_004236553.1) *Solanum tuberosum* (XP_006351473.1), *Nicotiana attenuata* (XP_019261423.1), *Nicotiana tabacum* (XP_016469144.1), *Oryza sativa* (XP_015615209.1), *Helianthus annuus* (XP_021972959.1), *Sesamum indicum* (XP_011083758.1). The phylogenetic trees were constructed using Phylogeny.fr (http://www.phylogeny.fr/) by bootstrap method. The scale bar indicates the branch lengths. **(B)** Alignment of AoCYP94B1 derived amino acid sequence with AoCYP94B1 of other crop species using MultAlin. The Cytochrome P450 cysteine heme-iron ligand signature motif, highlighted in black square, was well conserved.

**Supplemental Fig. 2: *pAtCYP94B1::GUS* expression and growth of 4-week-old WT, *atcyp94b1* and *35S::AoCYP94B1 Arabidopsis* plants:** (**A**) *pAtWRKY33::GUS* expression analysis in one-week-old seedling roots upon 50 mM NaCl treatment. Scale bar=500 µm. **(B)** Growth of the one-month-old, soil grown WT, *atcyp94b1* mutant and three independent lines of *35S::AoCYP94B1* heterologously expressed in the mutant background plants shown under untreated condition, under salt-treated condition and after recovery growth in normal water for one week. Scale bar=10 mm. wk; week.

**Supplemental Fig. 3: WT, *atcyp94b1*, *35S::AoCYP94B1* and *AtCYP94B1::AtCYP94B1* complementation lines of *Arabidopsis* responded similarly to mannitol treatment:** (**A**) Comparison of seedling growth among WT, *atcyp94b1* mutant and three independent lines of *35S::AoCYP94B1* heterologously expressed in the mutant background. (**B**) Growth of WT, *atcyp94b1* and *AtCYP94B1::AtCYP94B1* complementation lines. Surface sterilized and cold stratified seeds were sown on MS agar plates with or without mannitol (50 and 75 mM). Photographs were taken at the end of one week after germination. Scale bar=10 mm.

**Supplemental Fig. 4: FDA penetration into endodermal cells show functionality of suberin:** One-week-old WT, *atcyp94b1*, *pAtCYP94B1::AtCYP94B1* and *35S::AoCYP94B1* seedlings were incubated in FDA for 1 min, rinsed and images at the corresponding positions were taken using confocal microscopy. **(A, B)** FDA penetration into endodermal cells in the undifferentiated zone. **(C, D)** FDA penetration into endodermal cells in the non-suberized zone. Scale bar= 10 µm, ep: epidermis, co: cortex, en: endodermis. n=10 for all the analyses.

**Supplemental Fig. 5: Heterologous expression of *AoCYP94B1* increases deposition of apoplastic barrier (SL) in the roots of transgenic rice seedlings:** (**A**-**D**) Cross sections were made from the roots of 4-week-old hydroponically grown, salt-treated WT and *pUBI::AoCYP94B1* plants at varying lengths from the root tip: apical (0–20 mm), mid (20–50 mm) and basal (50–80 mm). For visualizing SL, sections were stained with Fluorol Yellow 088. Images of endodermal SL in the apical, mid and basal regions of (**A**) WT and (**B**) *pUBI::AoCYP94B1*_1. Images of exodermal SL in the apical, mid and basal regions of (**C**) WT and (**D**) *pUBI::AoCYP94B1*_1. Arrowheads indicate the presence of SL. Asterisks indicate endodermal cells lacking SL. Images were taken from sections made using at least three seedlings. Scale bar= 20 µm. en: endodermis, ex: exodermis.

**Supplemental Fig. 6: Supplemental Fig. 6: Promoter analysis of *AtCYP94B1*: (A)** *AtCYP94B1* promoter fragment showing stress-related transcription factor binding domains such as WRKY (red), MYB (pink) and MYC (pale pink). The DREME/MEME software suite (Bailey, 2011) was used to perform stringent motif searches within a 2000-bp region upstream of start codon of the coding region of *AtCYP94B1*. (**B**) *AtCYP94B1* promoter fragment used for Luciferase assay. Two WRKY binding domains highlighted in yellow were mutated by site directed mutagenesis (TTGAC to TTacC).

**Supplemental Fig. 7: *A. officinalis* WRKY33 shares high sequence similarity with *Arabidopsis* WRKY33:** A phylogenetic tree was constructed using the deduced amino acid sequences of *AoWRKY33* along with *Theobroma cacao* (XP_017977471.1), *Populus trichocarpa* (XP_002323637.2), *Ricinus communis* (EEF35722.1), *Jatropha curcas* (XP_012089749.1), *Solanum lycopersicum* (XP_004246308.1), *Nicotiana tabacum* (NP_001311970.1), *Capsicum chinense* (PHU06953.1), *Capsicum annuum* (NP_001311528.1), *Sesamum indicum* (XP_020553388.1), *Coffea arabica* (XP_027068692.1), *Coffea eugenioides* (XP_027173412.1), *Arabidopsis thaliana* (NP_181381.2). The phylogenetic trees were constructed using Phylogeny.fr (http://www.phylogeny.fr/) by bootstrap method. The scale bar indicates the branch lengths. (B) Sequence alignment of the derived amino acid sequences of *AoWRKY33* with *Arabidopsis AtWRKY33*. WRKY33s have two WRKY domains. Conserved WRKY domains are highlighted in yellow and conserved sequences are shown in red.

**Supplemental Fig. 8: Tissue specific expression of *pAtWRKY33::GUS*:** Expression of *pAtCYP94B1::GUS* in various tissues of mature plants bearing siliques. Scale bar= 500 µm.

**Supplemental Fig. 9: Details of *atcyp94b1* mutant and heterologous expression lines:** (**A**) Genetic map of *atcyp94b1* T-DNA insertion used in generating *35S::AoCYP94B1* lines. (**B**) Genotyping PCR confirms the homozygosity of the *atcyp94b1* mutant lines. (**C**) qRT-PCR shows suppressed expression of *AtCYP94B1* in *atcyp94b1* T-DNA insertional mutants. (**D**) qRT-PCR shows high level of expression of *35S::AoCYP94B1* in the *atcyp94b1* mutant background. * indicates the lines used for further analysis. Relative expression levels of transcripts with reference to *Ubiquitin 10* transcript levels, qRT-PCR data represent means ± SD from 3 biological replicates each with 3 technical replicates.

**Supplemental Fig. 10: Details of *Arabidopsis AtWRKY*33 mutant:** (**A**) Genetic map of *atwrky33* T-DNA insertion used. (**B**) Genotyping PCR confirms the homozygosity of the *atwrky33* mutant lines. (**C**) qRT-PCR shows suppressed expression of the *AtWRKY33* gene in *atwrky33* T-DNA insertional mutants. Relative expression levels of transcripts with reference to *Ubiquitin 10* transcript levels, qRT-PCR data represent means ± SD from 3 biological replicates each with 3 technical replicates.

